# Feedback regulation of BMP signaling by *C. elegans* cuticle collagens

**DOI:** 10.1101/686592

**Authors:** Uday Madaan, Lionel Faure, Albar Chowdhury, Shahrear Ahmed, Emma J. Ciccarelli, Tina L. Gumienny, Cathy Savage-Dunn

**Author notes:** Corresponding author: Cathy Savage-Dunn, Department of Biology and PhD Program in Biology, Queens College and the Graduate Center, CUNY, 65-30 Kissena Blvd., Flushing, NY 11367. these authors contributed equally (Uday Madaan, Lionel Faure).

## Abstract

Cellular responsiveness to environmental cues, including changes in extracellular matrix (ECM), is critical for normal processes such as development and wound healing, but can go awry, as in oncogenesis and fibrosis. One type of molecular pathway allowing this responsiveness is the bone morphogenetic protein (BMP) signaling pathway. Due to their broad and potent functions, BMPs and their signaling pathways are highly regulated at multiple levels. In *Caenorhabditis elegans*, the BMP ligand DBL-1 is a major regulator of body size. We have previously shown that DBL-1/BMP signaling determines body size through transcriptional regulation of cuticle collagen genes. We have now obtained evidence of feedback regulation of DBL-1/BMP by collagen genes. We analyzed four DBL-1-regulated collagen genes that affect body size. Here we show that inactivation of any one of these cuticle collagen genes reduces DBL-1/BMP signaling, as measured by a Smad activity reporter. Furthermore, we find that depletion of these collagens reduces GFP::DBL-1 fluorescence, and acts unexpectedly at the level of *dbl-1* transcription. We therefore conclude that cuticle, a type of ECM, impinges on DBL-1/BMP expression and signaling. In contrast to other characterized examples, however, the feedback regulation of DBL-1/BMP signaling by collagens is likely to be contact-independent, due to the physical separation of the cuticle from DBL-1-expressing cells in the ventral nerve cord. Our results provide an entry point into a novel mechanism of regulation of BMP signaling, with broader implications for mechanical regulation of gene expression in general.

## Introduction

Cells are exquisitely sensitive to their environments, to which they not only respond but also create and maintain in the secretion of extracellular matrix (ECM). Cells maintain the homeostasis of their ECM by regulating the production of secreted collagens, other ECM components, and ECM-modifying enzymes. Loss of this regulation can result in cell death or fibrosis. One of the regulators of ECM homeostasis is the Transforming Growth Factor beta (TGF-β) family signaling pathway. TGF-β signaling is responsive to changes in matrix stiffness, is regulated by collagens, and in turn regulates collagen synthesis [1, 2]. However, *in vitro* studies of this interaction are limited in their ability to recapitulate the complexities of cell-matrix interactions, much less the indirect interactions of cells within organs and ECM, and *in vivo* systems have thus far been limited.

We have established *C. elegans* as an *in vivo* model to study the interplay between TGF-β signaling and ECM. The *C. elegans* ECM includes both basement membrane, which lines internal organs, and cuticle, which is secreted apically from hypodermal tissue to protect the organism and is shed at each molt [3-5]. The cuticle is composed of multiple collagen layers coated by a lipid-rich epicuticle [6, 7]. Cuticle collagens are encoded by a large multi-gene family with over 170 members. Some cuticle collagens are expressed in each developmental stage in which a cuticle is synthesized, while others are expressed stage-specifically [8]. Accordingly, the cuticle at each developmental stage contains different specific collagens, and is organized into distinct structures. For example, the lateral ridges known as alae are only present on L1, dauer, and adult cuticles [5].

The TGF-β family includes Bone Morphogenetic Proteins (BMPs), highly conserved members such as vertebrate BMP2, Drosophila Dpp, and *C. elegans* DBL-1 peptides. The DBL-1 ligand is not only highly conserved at the sequence level, but it signals through a conserved, canonical signaling pathway as well. This pathway includes the type I and type II receptors and intracellular Smad signal transducers [9]. The DBL-1 ligand is secreted by ventral cord neurons that are nestled in the basal side of the hypodermis [10]. Receptors and Smad signal transducers for DBL-1 are present in the hypodermis, which receives the DBL-1 signal [11, 12]. The DBL-1 signaling pathway has been shown to function in a number of developmental and homeostatic functions, but is not essential for viability. There is growing evidence that DBL-1 signaling modulates cuticle structure and function. For example, the DBL-1 pathway plays a major role in regulating body size and morphology, permeability to drugs, and resistance to infection, traits associated with cuticle function [13]. Loss of *dbl-1* causes physical changes in cuticle morphology [13]. Finally, genes encoding cuticle collagens and other ECM constituents and enzymes are known transcriptional targets of the DBL-1 pathway [14-16].

We previously characterized four of these DBL-1-regulated collagen genes, *col-41, rol-6, col-141*, and *col-142*, and demonstrated that they are involved in body size regulation [17]. Here we report that these four DBL-1-regulated collagens reciprocally modulate the DBL-1 pathway. Using three different activity reporters, we show that depletion of individual collagen genes impacts the DBL-1 pathway at the levels of Smad activity, DBL-1 protein, and *dbl-1* expression. We propose a model in which the presence of these cuticle collagens modifies properties of the cuticle to which DBL-1-expressing cells are sensitive.

## Materials and Methods

### Strains

*C. elegans* strains were grown at 20°C using standard methods [18]. All experiments were performed at 20°C. In addition to strains generated in this work, the following strains were used: LW2436 *jjIs2436[pCXT51(RAD-SMAD)* + *LiuFD61(mec-7p::mRFP)]*, TLG281 *rrf-3(pk1426)*; *texIs100[dbl-1p::GFP::dbl-1* + *ttx-3p::mRFP]*; *dbl-1(nk3)*, BW1935 *unc-119(ed3); ctIs43[dbl-1p::GFP* + *unc-119(*+*)]; him-5(e1490)*, MT2709 *rol-6(e187n1270)*, CS678 *col-141(qc25[2xNLS::GFP])*, CS637 *qcEx131[col-141(*+*) col-142(*+) + *myo-2p::GFP].*

### RNA interference (RNAi)

RNAi by feeding was performed as described [19-21]. RNAi clones that target the unique (non-Gly-X-Y repeat) regions of collagen genes were generated by cloning PCR fragments encoding these regions into L4440 vector and transforming bacterial strain HT115 [17].

### RAD-SMAD reporter and *dbl-1* transcriptional reporter imaging

Fluorescence images were taken at 40x using a Zeiss Apotome microscope. For fluorescence quantification, images were taken at 40x. Fluorescence intensity was determined by manually selecting each individual nucleus and quantifying intensity using ImageJ software. Empty vector (L4440) RNAi was used as a control. In addition to the parent reporter strains, crosses were performed to generate CS657 *ctIs43;rol-6(e187n1270)*, CS675 *ctIs43;col-141(qc25)*, and CS677 *ctIs43;qcEx131.*

### Body size measurements and GFP::DBL-1 imaging

Populations of TLG281 *rrf-3(pk1426)*; *dbl-1(nk3)*; *texIs100* were staged by a standard bleaching technique and grown overnight at 20°C in M9 to hatch and arrest at the L1 stage [22]. Starved hatchlings were plated on bacteria expressing double-stranded RNAi against each gene of interest and grown at 20°C for five days, until all animals were young adults. RNAi against the unrelated gene C06C3.5 was used as a control. In addition to the RNAi treatments, the *rol-6(e187n1270)* allele was crossed with *texIs100* to generate CS636 *rol-6(e187n1270); texIs100.*

Adult animals were picked up and were immobilized with 0.2mM levamisole in M9 buffer on a 10% agarose pad before mounting [23]. Confocal images were acquired using either a Nikon A1R confocal system or Nikon Eclipse Ti-E confocal system. We used a 60×/1.40 NA oil CFI Plan Apo VC objective and the NIS-Elements Advanced Research software (Nikon Instruments, Inc., Melville, NY, USA) to measure and quantify the intensity of the GFP::DBL-1 protein under different conditions. The intensity of the GFP tagged protein for a selected area was calculated by calculating the mean intensity associated with this area minus the mean intensity of the background. Body length was measured using 10× magnification and NIS AR software.

## Results

### Depletion of DBL-1-regulated cuticle collagen gene products impacts DBL-1 pathway activity

To determine whether cuticle collagens impact DBL-1 pathway activity, we used an artificial transcriptional reporter containing multiple Smad binding sites driving expression of GFP (RAD-SMAD reporter [24]). We used RNAi to deplete expression of four cuticle collagen genes, *col-41, rol-6, col-141*, and *col-142*, in animals carrying this RAD-SMAD reporter. RNAi was performed with dsRNA targeting the gene-specific amino-termini that do not encode Gly-X-Y repeats. At the L4 stage, depletion of *rol-6* caused significant reduction in RAD-SMAD activity in the hypodermis (Fig. 1A). Compared with control animals fed bacteria carrying the empty vector, the depletion of *col-141* or *col-142* reduced RAD-SMAD activity in L4 animals. The reduction in activity was statistically significant in some but not all trials. In one-day old young adults, a more pronounced reduction in fluorescence was seen after *rol-6, col-141*, and *col-142* depletion (Fig. 1B). At both L4 and adult stages, RNAi targeting *col-41* led to a large variance in fluorescence intensity between individual nuclei, with many nuclei showing reduced expression (Fig. 1A, B). We conclude that DBL-1 Smad signaling in the hypodermis is somehow impacted by the collagen composition of the cuticle.

**Figure 1.**
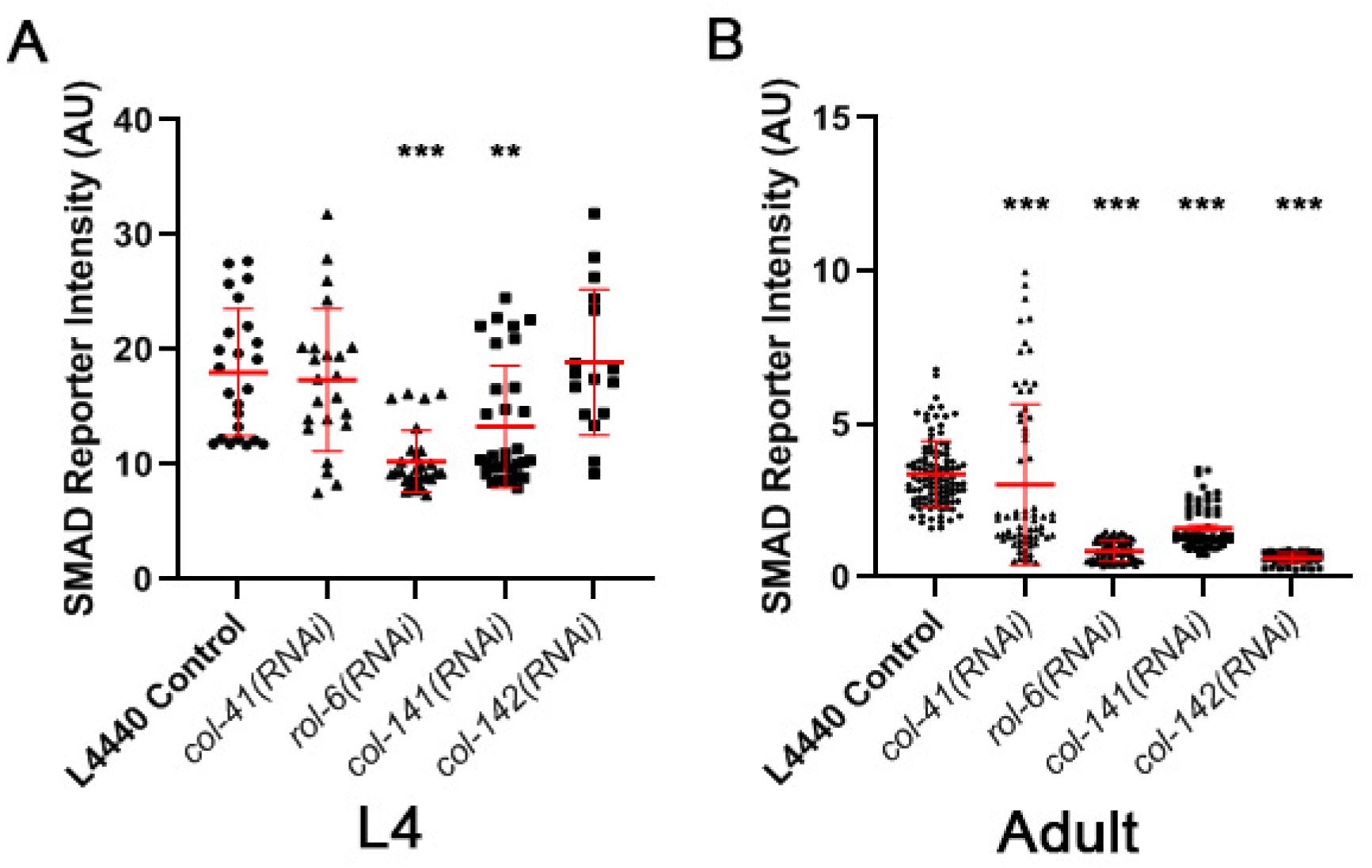
Loss of cuticle collagen gene function reduces activity of the DBL-1/BMP pathway. A. L4 animals. B. Young adults. RNAi was used to deplete collagen gene function. Pathway activity was assayed using a reporter containing Smad binding sites. AU: arbitrary units. Bars show mean and standard deviation of the data. Statistical significance compared with control was determined by Brown-Forsythe and Welch ANOVA tests. * p < 0.05; ** p < 0.01; *** p < 0.001.

### Manipulation of DBL-1-regulated cuticle collagen genes suppresses long body size phenotype of *dbl-1* overexpression

We previously showed that cuticle collagen genes are effectors of DBL-1 regulation of body size [17]. In the previous study, using targeted RNAi, overexpression strains, and loss-of-function mutants, we found that *col-41* is a positive regulator, *rol-6* is a dose-dependent regulator, and *col-141* is a negative regulator of body size [17]. To elucidate further how these cuticle collagens interact with the DBL-1 pathway in body length regulation, we analyzed their body size phenotypes in a strain (TLG281) containing an integrated transgene (*dbl-1(oe)*) that overexpresses functional GFP-tagged DBL-1, which results in a long body size. First, we examined the interaction of TLG281 (*dbl-1(oe)*) with RNAi targeting *col-41* or *rol-6*, which cause small body size phenotypes in the wild type. If these genes act downstream of *dbl-1* for body size regulation, we expect their depletion to suppress the long body size phenotype of *dbl-1(oe). dbl-1(oe)* animals fed on bacteria expressing *col-41(RNAi)* showed a significant reduction of their long body size in adult animals compared to our control (C06C3.5 pseudogene RNAi) (Fig. 2B). We also observed that feeding *rol-6(RNAi)* caused a significant reduction in the long body size phenotype of *dbl-1(oe)* in L4 and adult animals (Fig. 2A, B). Finally, we examined *rol-6(e187n1270)*, a loss-of-function mutant (hereafter *rol-6(lf))* that results in reduced body length in wild-type animals [17, 25]. In CS636 (*rol-6(lf)* expressing GFP::DBL-1), we observed a significant reduction of the long body size phenotype compared to *dbl-1(oe)* animals for both L4 and adult stages (Fig. 2A, B). Together, these results support our hypothesis that *col-41* and *rol-6* act downstream of DBL-1 to regulate body size.

**Figure 2.**
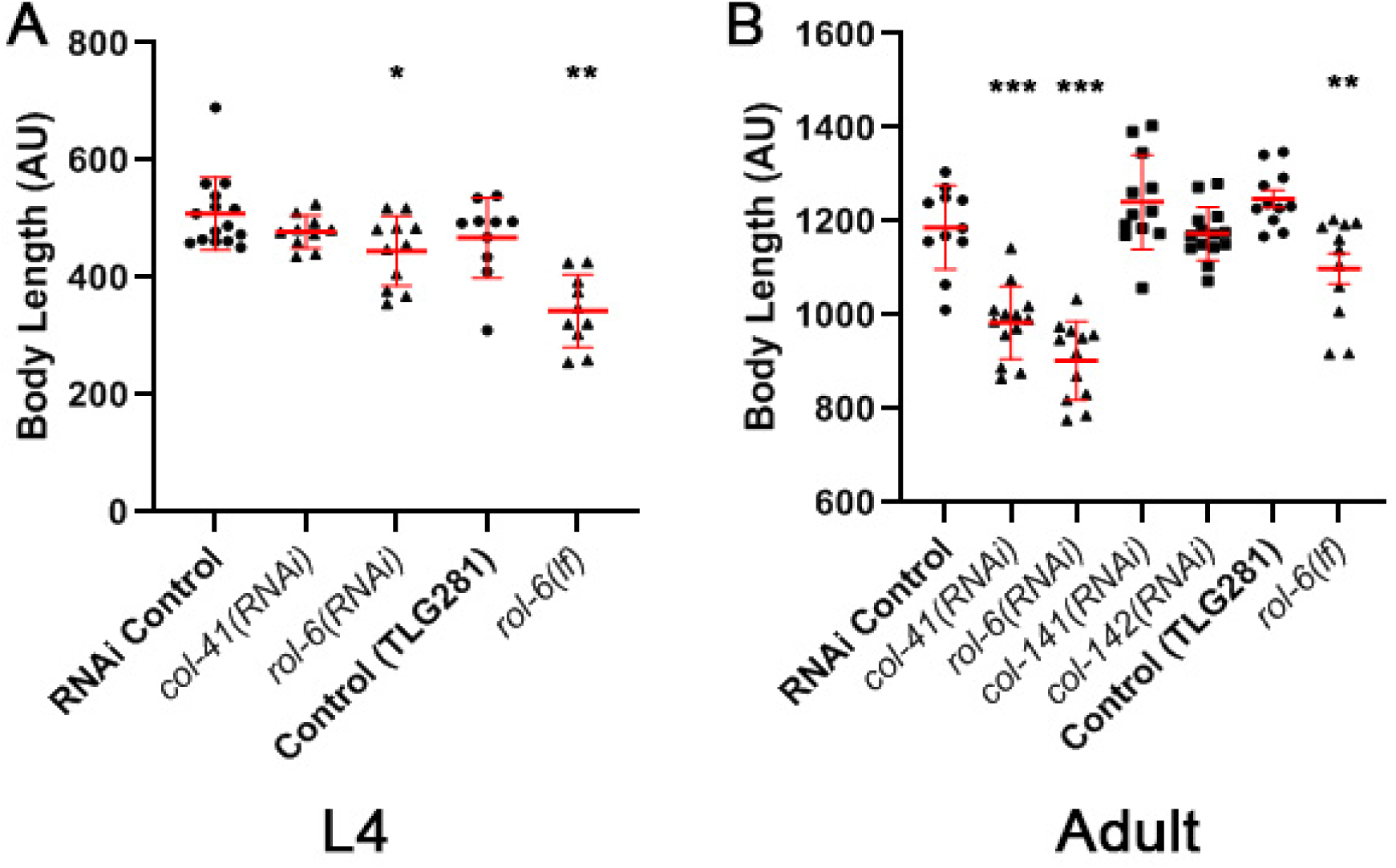
Genetic interactions between cuticle collagen genes and DBL-1 overexpression in body size regulation. A. L4 animals. B. Young adults. A strain carrying *texIs100*, a multi-copy array that overexpresses GFP::DBL-1 causing a long body size phenotype, was treated with collagen RNAi or crossed with *rol-6(lf)* as indicated. Body length was measured in one-day old adults. AU: arbitrary units. Bars show mean and standard deviation of the data. Statistical significance compared with control was determined by Brown-Forsythe and Welch ANOVA tests. * p < 0.05; ** p < 0.01; *** p < 0.001.

In a previous study, we showed that *col-141(RNAi)* causes an increase in body size, while overexpression of *col-141* and *col-142* reduces body size in wild-type animals [17], suggesting that *col-141* and *col-142* are negative regulators of body length in *C. elegans*. We tested for genetic interactions between *dbl-1(oe)* and *col-141(RNAi)* or *col-142(RNAi).* If these genes act independently in body size regulation, then we would expect additive effects, with RNAi of *col-141* or *col-142* further increasing the long body size of *dbl-1(oe).* When *col-141* or *col-142* was depleted by RNAi in the *dbl-1(oe)* background, no further increase in body size was seen (Fig. 2B). This result is consistent with these genes acting in the same pathway as DBL-1 to regulate body size. Combined with our previously published data [17] and the results from the RAD-SMAD reporter (Fig. 1), these new results (Fig. 2) provide evidence of bidirectional signaling between the DBL-1 pathway and cuticle collagen genes.

### Intensity of GFP::DBL-1 is reduced upon loss of DBL-1-regulated collagens

The effect of collagen gene manipulation on the RAD-SMAD transcriptional reporter could occur either at the level of the ligand, or on downstream components of the signaling pathway, or possibly both. To distinguish between these possibilities, we first asked whether collagen gene manipulation alters DBL-1 protein distribution. We used the strain expressing functional GFP::DBL-1 described above [13] fed on bacteria transformed with different RNAi targets and quantified protein fluorescence. We found that depletion of any of the collagen genes *col-41, rol-6, col-141*, or *col-142* by RNAi reduces the intensity of GFP::DBL-1 fluorescence in adults compared to our control (C06C3.5 pseudogene RNAi) (Fig. 3B), although the reduction upon *col-41(RNAi)* does not reach statistical significance (p = 0.26). These results support the data obtained with the RAD-SMAD experiment (Fig. 1). We also found that RNAi of *col-41*, but not of *rol-6*, caused a significant reduction in GFP intensity at the L4 stage (Fig. 3A). As in the RAD-SMAD experiment, RNAi of *col-41* caused a large variance in fluorescence intensity (Figs. 1 and 3). Finally, we verified these results by measuring the fluorescence intensity of GFP::DBL-1 in the *rol-6(lf);dbl-1(oe)* strain compared with *dbl-1(oe)*. In *rol-6(lf)* mutants, GFP::DBL-1 fluorescence was significantly reduced in L4 animals (Fig. 3A) and slightly, but not significantly, reduced in adults (Fig. 3B). Taken together we therefore conclude that loss of these cuticle collagen isoforms leads to reduction in DBL-1 protein levels. The reduced RAD-SMAD activity observed in Fig. 1 is therefore likely due, at least in part, to changes in DBL-1 accumulation.

**Figure 3.**
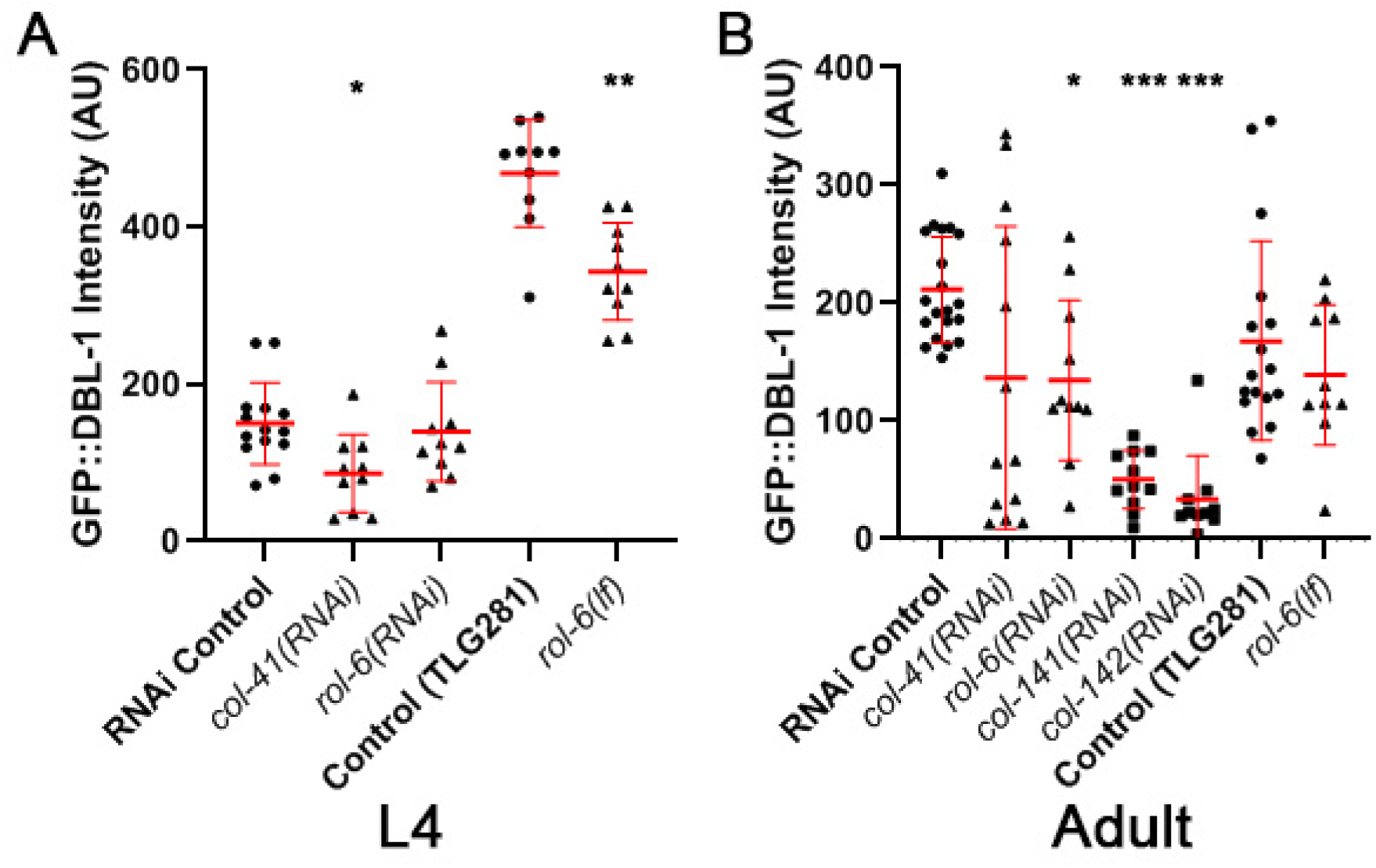
GFP::DBL-1 accumulation responds to manipulation of cuticle collagen genes. A. L4 animals. B. Young adults. A strain carrying *texIs100* that expresses functional GFP tagged DBL-1 was treated with collagen RNAi or crossed with *rol-6(lf)* as indicated. Fluorescence intensity was measured in one-day old adults. AU: arbitrary units. Bars show mean and standard deviation of the data. Statistical significance compared with control was determined by Brown-Forsythe and Welch ANOVA tests. * p < 0.05; ** p < 0.01; *** p < 0.001.

### DBL-1-regulated collagens affect expression of a *dbl-1* transcriptional reporter

We observed that the genetic manipulation of cuticle collagens caused a reduction in GFP::DBL-1 protein levels (Figs. 2 and 3), but these experiments do not distinguish the level at which such regulation occurs. We considered two models: regulation of DBL-1 protein distribution or stability and regulation of *dbl-1* transcription. To distinguish between these possibilities, we used a transcriptional reporter in which GFP is driven by the *dbl-1* promoter [10]. We subjected this strain (BW1935) to RNAi depletion of DBL-1-regulated collagen genes. Intriguingly, we found that loss of these cuticle components is associated with a significant reduction of the expression of the *dbl-1* transcriptional reporter at both the L4 and adult stages for all conditions tested (Fig. 4). The large variability in Smad reporter activity and GFP::DBL-1 levels following *col-41(RNAi)* was not seen for the *dbl-1* transcriptional reporter. These results demonstrate that cuticle collagen genes are capable of regulating DBL-1 at the level of transcription, an unexpected finding due to the physical separation of DBL-1-expressing neurons from the cuticle.

**Figure 4.**
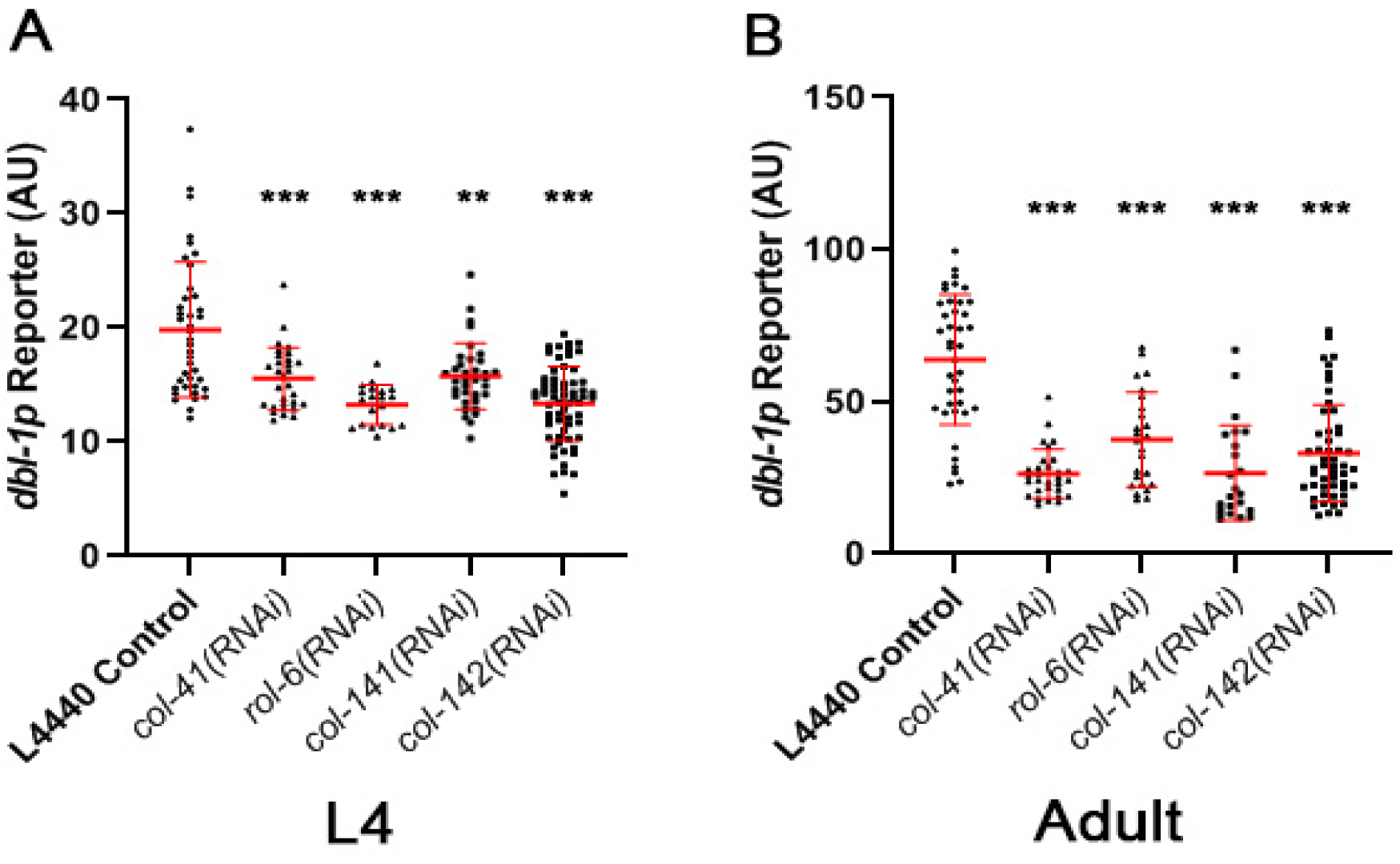
Loss of cuticle collagen gene function reduces expression of a DBL-1 transcriptional reporter. A. L4 animals. B. Young adults. A strain carrying *ctIs43*, a transgene expressing GFP driven by the *dbl-1* promoter, was treated with collagen RNAi as indicated. AU: arbitrary units. Bars show mean and standard deviation of the data. Statistical significance compared with control was determined by Brown-Forsythe and Welch ANOVA tests. * p < 0.05; ** p < 0.01; *** p < 0.001.

Finally, we crossed the *dbl-1* transcriptional reporter into three collagen mutant strains: *rol-6(lf), col-141 col-142(oe)*, and a *col-141(lf)* deletion allele generated by CRISPR/Cas9 genome editing [17]. We quantified the GFP intensity from *dbl-1p::gfp* in young adults as a measure of *dbl-1* transcription levels. Surprisingly, in *rol-6(lf)* mutants, GFP expression became undetectable (Fig. 5B). Thus, loss of *rol-6* has a major effect on *dbl-1* expression. Similarly, *col-141(lf)* mutants have slightly reduced *dbl-1* expression (Fig. 5C, E). In contrast, *col-141 col-142(oe)* animals have normal levels of *dbl-1* reporter expression (Fig. 5D, E). These results from the *rol-6(lf)* and *col-141(lf)* mutant strains corroborate the findings from the respective RNAi experiments. We conclude that cuticle collagens participate in feedback regulation of DBL-1.

**Figure 5.**
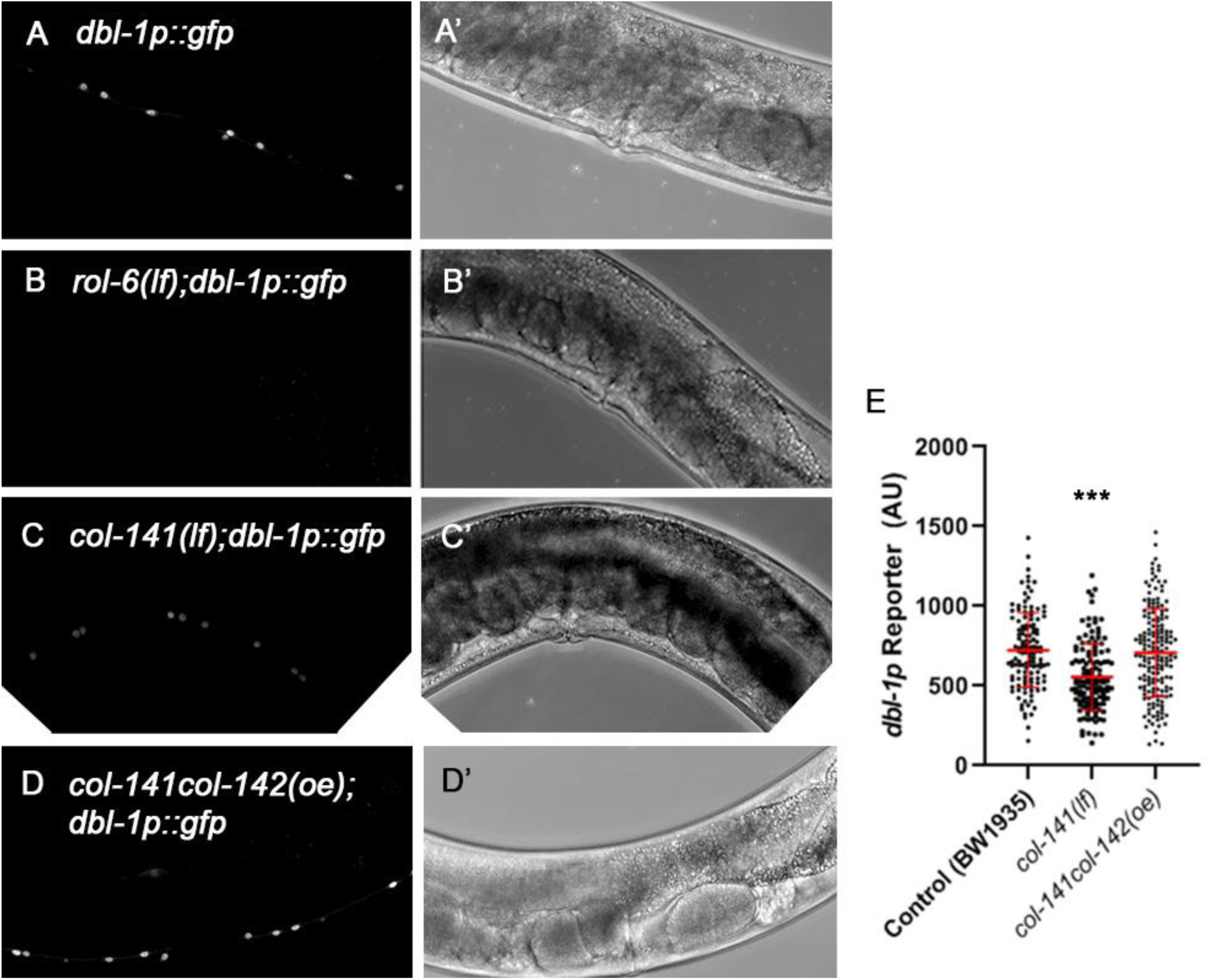
DBL-1 transcriptional reporter expression in collagen mutant and overexpression strains. The transgene *ctIs43*, expressing GFP driven by the *dbl-1* promoter, was visualized in **A.** wild-type, **B.** *rol-6(lf)*, **C.** *col-141(lf)*, and **D.** *col-141 col-142(oe)* backgrounds. **A-D.** Representative GFP fluorescence images. **A’-D’.** The same animals visualized with Nomarski DIC optics. Ventral is downward. **E.** Fluorescence intensity was measured in one-day old adults. AU: arbitrary units. Bars show mean and standard deviation of the data. Statistical significance compared with control was determined by Brown-Forsythe and Welch ANOVA tests. * p < 0.05; ** p < 0.01; *** p < 0.001.

## Discussion

In this work, we investigated the reciprocal interactions between the DBL-1/BMP pathway and specific cuticle collagens, COL-41, ROL-6, COL-141, and COL-142. In previous work, we showed that *col-41, rol-6, col-141*, and *col-142* are transcriptional targets of DBL-1 signaling [17]. They are positive, negative, or dose-sensitive regulators of body size [17]. Moreover, DBL-1 signaling mutants have altered cuticle morphology and function [13]. Here we extended those studies to provide evidence that these cuticle collagen genes act downstream of DBL-1 in body size regulation. We used three different measures of DBL-1 pathway signaling to demonstrate that alterations of these cuticle collagen genes, in turn, feed back to modulate DBL-1 signaling. Interestingly, this postulated feedback regulation is consistent with a previous study that shows the ADAMTS secreted metalloprotease ADT-2 modulates both cuticle collagen organization and DBL-1 signaling activity [26].

These four DBL-1-regulated collagen genes can be divided into two groups based on temporal expression patterns and loss-of-function phenotypes. One group, consisting of *col-41* and *rol-6*, each has a peak of expression in the second larval (L2) stage, and loss of these genes causes a mild small body size phenotype. DBL-1 signaling activates expression of these genes at the L2 stage and represses their expression in adults [17]. Loss of *col-41* caused reduction in *dbl-1* expression, GFP::DBL-1 protein fluorescence, and Smad reporter activity (Figs. 1, 3, and 4). Strikingly, *col-41(RNAi)* consistently resulted in a large variation in Smad activity reporter and GFP::DBL-1 levels, and in some experiments this may have contributed to a lack of statistical significance. This variance was not seen in the level of the *dbl-1* transcriptional reporter, which suggests the existence of a post-transcriptional compensatory mechanism for DBL-1 levels under these conditions. This intriguing observation invites further elucidation.

Loss of *rol-6* had the most profound effects on DBL-1 signaling, causing significant reductions in expression of a Smad activity reporter, GFP::DBL-1 fusion protein, and a *dbl-1* transcriptional reporter. The effects of *rol-6* on body size are also substantial. Reduction of the *rol-6* transcript by *rol-6(RNAi)* or by a *rol-6(lf)* mutation both cause a small body size [17] and significantly suppress the long body size of *dbl-1(oe)* (Fig. 2). Furthermore, *rol-6(oe)* is also small, indicating a dose-sensitivity for this gene product in body size regulation. These results extend ROL-6’s functions to include regulation of DBL-1 signaling.

The second group comprises *col-141* and *col-142*, which have a peak of expression in the adult stage. Loss of *col-141* causes a mild long body size phenotype. DBL-1 Smads directly regulate these genes, repressing expression at the L2 stage and activating their expression in adults [17]. In spite of the opposite functions of *col-41* and *rol-6* in body size determination, *col-141* and *col-142* act similarly to those genes in reducing DBL-1 signaling activity, as seen in three independent activity reporters (Figs. 1, 3, and 4). Like *col-141(RNAi)* (Fig. 4), *col-141(lf)* reduced *dbl-1* reporter transcription (Fig. 5). Conversely, *col-141 col-142(oe)*, which leads to small animals, does not affect the *dbl-1* transcriptional reporter. These results dissociate the collagens’ effects on body length from DBL-1 pathway activity, so the effect of collagen loss on DBL-1 signaling likely depends on other properties of the cuticle in these backgrounds.

It is well established in various contexts that members of the TGF-β family, including BMPs, regulate ECM, with important consequences for biological function, which we can refer to as “forward” regulation of the ECM [1, 27]. Moreover, the ECM can participate in “reverse” or “feedback” regulation of TGF-β or BMP signaling. For example, type IV collagen controls BMP signaling in Drosophila by direct binding of Dpp [28-30]. In addition, the ECM protein fibrillin directly binds TGFβs and BMPs [31]. Mutations in FBN1 result in Marfan syndrome, for which some of the pathologies are attributed to increased TGF-β bioavailability [32]. Similarly, the interaction between fibrillin1 and BMP4 regulates BMP4 bioavailability [33]. Each of these interactions occurs via direct physical contact between ligands and the ECM. Our work, in contrast, describes a novel role for contact-independent regulation of BMP ligand expression and activity by the ECM. Fig. 6 illustrates our model, including the locations of relevant structures. The cuticle forms the outermost barrier. Its components are synthesized by the underlying hypodermis and secreted to the apical (outer) surface. DBL-1 is secreted from neurons apposed to the basal (internal) side of the hypodermis. DBL-1-expressing cells are thus physically separated from the cuticle, making physical interactions between these cells and cuticle collagens unlikely.

**Figure 6.**
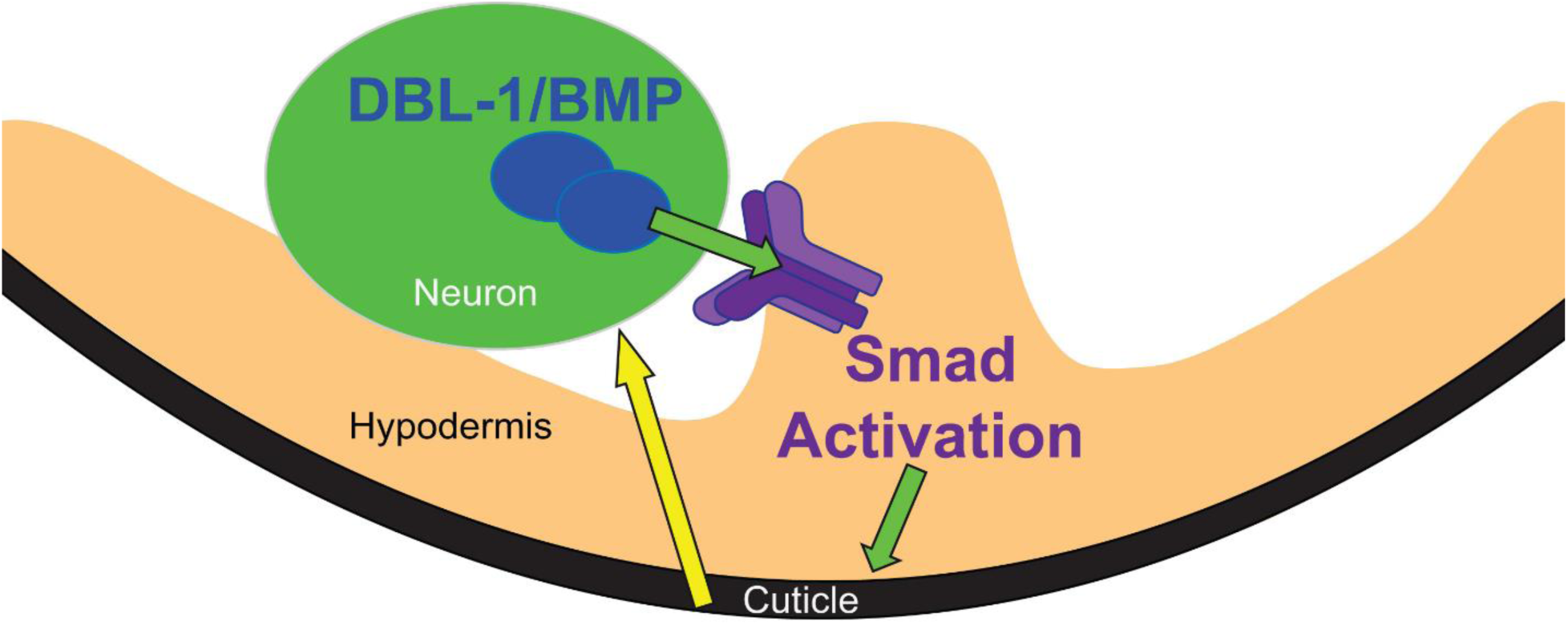
A model for DBL-1/BMP pathway regulation by cuticle collagens. Green arrows represent forward DBL-1 signaling that regulates cuticle collagen expression and body size. DBL-1/BMP is expressed by and secreted from neurons of the ventral cord. It binds to receptors on the hypodermis and activates Smad signaling. Smad activation causes changes in gene expression leading to altered collagen composition of the cuticle. Yellow arrow depicts feedback signaling from the cuticle to DBL-1/BMP expression.

How can a contact-independent mechanism operate? Biomechanical properties of a tissue can influence gene expression. For example, mesenchymal stem cells commit to different differentiated lineages depending on matrix stiffness [34]. In another example, compression of Drosophila embryos (which may mimic forces generated during morphogenetic movements) induces ectopic expression of the transcription factor Twist [35]. Alterations of the *C. elegans* cuticle have also been shown to cause changes in signaling pathways in non-hypodermal tissues. For example, loss of some cuticle collagens alters widely expressed SKN-1/Nrf transcription factor activity and stress responses [36]. Interestingly, classical work on the GLP-1/Notch receptor in *C. elegans* identified mutations in seven cuticle collagen genes that suppress the pharyngeal and germline phenotypes of partial-loss-of-function *glp-1* alleles [37]. They also found the severity of the body size defect associated with loss of collagen function did not correlate with *glp-1* suppression. In this work, we show a similar effect on BMP signaling by loss of a different set of cuticle collagens. The reduction of DBL1/BMP expression does not correlate with the alteration in body size in these collagen knock-downs. Specifically, RNAi inhibition of *col-41* or *rol-6* results in small animals, while RNAi against *col-141* results in long animals, but each of these treatments reduces DBL-1 reporter activity. Therefore, it is not the animal’s size but some other feature of the altered cuticle that is responsible for the feedback regulation.

Our model (Fig. 6) summarizes these observations. The neurons expressing DBL-1 may experience biomechanical forces, such as tension or compression, which modulate gene expression, including *dbl-1* expression. Alterations in the collagen composition of the cuticle affect these forces, leading to changes in gene expression. This phenomenon may be a more widespread mechanism of contact-independent regulation of gene expression and cell signaling, for which the *C. elegans* cuticle may be a useful model. Future studies will identify the biomechanical properties of the *C. elegans* cuticle responsible for this proposed feedback. In summary, the DBL-1/BMP pathway provides a unique *in vivo* model to study bidirectional interactions between cell signaling and the ECM in the context of the intact organism. We propose that reciprocal interactions permit robust yet environmentally-responsive control of body size.

## Acknowledgments

We thank Jun (Kelly) Liu for providing strains and helpful conversations. We thank Alicia Meléndez for helpful feedback on the manuscript. This work was supported in part by National Institutes of Health R15GM112147 to CSD and R01GM097591 and TWU departmental funds to TLG. Some strains were provided by the CGC, which is funded by NIH Office of Research Infrastructure Programs (P40OD010440). This work was carried out in partial fulfillment of the requirements for the Ph.D. degree from the Graduate Center of City University of New York (UM and EJC).

